# Skeletons in the Forest: Using Entropy-Based Rogue Detection on Bayesian Phylogenetic Tree Distributions

**DOI:** 10.1101/2024.09.25.615070

**Authors:** Jonathan Klawitter, Remco R. Bouckaert, Alexei J. Drummond

## Abstract

In a phylogenetic analysis, rogue taxa and clades are those that, due to their uncertain placement, contribute disproportionally to the variance in a set of phylogenetics trees. They can obscure clear evolutionary relationships and complicate the interpretation of results. While existing rogue detection algorithms focus on improving the consensus tree for a set of trees, we introduce algorithms based on phylogenetic entropy to improve the analysis in a Bayesian framework. In particular, we introduce a tool that extracts a skeleton distribution of the posterior tree distribution that contains the core relationships by removing a minimal subset of rogues. Additionally, we enable detailed analyses of rogues at both the taxon and clade levels, including the visualisation of different rogue placements.

## 1 Introduction

The phylogenetic pipeline consists of many steps from obtaining sequences, to constructing a multiple sequence alignment (MSA), selecting a model, running phylogenetic inference, and finally analysing the results. Errors and noise can arise from various sources such as sequencing errors, low-quality sequences, and misalignment resulting in noisy MSAs [12], model misspecification [11], and the stochasticity of the inference process. Given the automation of the pipeline and ever growing amounts of data used, we need algorithms that help detect and address these sources of noise. We focus here on the last stage of the pipeline where we have a set of phylogenetic trees.

Many phylogenetic inference methods produce a set of trees rather than a single, definitive tree, to capture the inherent uncertainty in the evolutionary relationships studied. This includes bootstrap samples from maximum likelihood analyses [14, 27, 32], maximum parsimony or distance based methods [15], and posterior tree distributions from Bayesian analyses [5, 17, 30]. A high variance among the trees in such a set directly impacts our ability to accurately identify evolutionary relationships and interpret the results of an analysis. Regardless of the original cause, the variance among the trees ultimately manifests as different placements of taxa and clades with respect to other clades. Taxa and clades with the most uncertain placement contribute the most to this variance and are called *rogues* (or *wandering taxa*), a term coined by Wilkinson [35]. Here we are interested in the problem of efficiently detecting rogues and obtaining a *skeleton* of the tree distribution (with rogues removed) that represents the highly supported evolutionary relationships.

A set of tree is often compacted into a *summary tree* for visualisation and analysis, where it is assumed that this *point estimate* is representative of the distribution over trees in the set [3]. A classical approach to obtain a point estimate for a set of trees from a bootstrap analysis is to construct the *majority rule consensus tree*, which contains all clades (splits, for unrooted trees) occurring in the majority of the trees [7]. There, rogue taxa cause a loss of resolution in the consensus tree, leading to polytomies where multiple taxa appear to diverge from a single node, rather than showing clear bifurcating relationships. The uncertainty introduced by rogue taxa can also lead to incorrect biological interpretations, such as the misplacement of taxa [34], or incorrect timing of divergence events. Errors in summary trees can affect post-hoc analyses such as ancestral traits reconstruction, comparative genomics, coevolution studies, or biogeographic reconstructions. The majority of rogue detection algorithms thus aim to improve both the resolution and the support of the clades (splits) in a consensus tree [1, 2, 13, 28, 31], which however is an NP-hard problem [9]. Though methods with a different focus exist [23, 33], and we refer to Fleming, Valero-Garcia, and Struck [12] for an overview on rogue detection algorithms. Furthermore, Wilkinson and Crotti [36] point out that particular methods might not be suitable for all types of data.

We approach the problem of rogue detection differently and instead of focusing on a summary tree, we consider rogues in the context of posterior (rooted) tree distributions from a Bayesian analysis. Here we introduce new algorithms for rogue detection and the construction of a skeleton tree distribution. To this end, we utilize *conditional clade distributions (CCDs)*, which offer better estimates of the true posterior tree distribution than the tree sample [3, 16, 22]. CCDs allow us to efficiently work with large sets of trees via the underlying CCD graph, which captures all clade and clade split relationships, instead of having to process all trees. In particular, with a CCD we can compute the *phylogenetic entropy* of the posterior tree distribution [25], which quantifies the uncertainty in the distribution in an information theoretic way. As the basis of our rogue detection, we use the difference in entropy of a CCD with and without a particular clade. This is similar to the information theoretic method proposed by Smith [31], which however only scores splits (working with unrooted trees) independently and focuses on improving the consensus tree. We demonstrate how our rogue detection algorithm can be used to analyse all clades of a posterior as well as used successively to unearth the skeleton underlying the posterior. Importantly, using a small well-calibrated simulation study with a popular model, we observe that the construction of the skeleton does not introduce a detectable bias in the estimated evolutionary model parameters.

An open source implementation is freely available in the CCD package for BEAST2^1^, which offers different tools for rogue detection and obtaining skeletons with different strategies and termination criteria. It is also integrated with DensiTree [6] allowing interactive rogue identification while displaying the resulting tree set.

## 2 Methods

In this section, we first give a description of CCDs, then define phylogenetic entropy, and describe how we use it in our rogue detection algorithms. We then outline some example applications and list the datasets we use in our experiments. Throughout this section, we use the term *tree* for a rooted binary phylogenetic tree without branch lengths, what is commonly referred to as the (rooted) tree topology. Furthermore, we consider a taxon also to be a (trivial) clade. We use the notation [*k*] for the set {1, …, *k*}.

### 2.1 Conditional Clade Distributions

A *CCD* is a tree distribution that parametrises tree space and offers an estimate of a posterior distribution of trees. While a CCD can model the whole tree space, it is in general used to only model the tiny fraction of tree space that corresponds to nearly 100% of a true posterior distribution.

There are three progressively more complex models, *CCD0, CCD1*, and *CCD2*, with *CCD0* making the strongest independence assumptions about how clades in one part of a tree influence clades in another part, and *CCD2* the weakest. (We describe the different models below.) As such, a CCD has a bias-variance trade-off for estimating the true posterior distribution. Namely, the simpler model CCD0 with stronger independence assumptions has a higher bias but lower variance and it is easier to populate its parameters. On the other hand, the more complex models CCD1 and CCD2 have a lower bias but a higher variance. In comparison, a sample of trees is another model of the posterior distribution where each tree is a parameter. While this model offers potentially a perfect fit, not all trees with non-negligible probability can be sampled in practice due the super-exponential growth of treespace. In our recent study [3], we provided evidence of this trade-off, for both point estimates and estimation of the posterior probability of trees. We found that for non-trivial inference problems, in order to estimate the parameters of a CCD1 (or even a CCD2) to a degree that a CCD1 offers a better estimate than a CCD0 requires a huge number of sampled trees.

To explain how a CCD represents a tree distribution, we first need to describe its structure and the conditional clade probability of a clade split. First, underlying a CCD *D* is a *CCD graph G*, which is a directed bipartite graph with the two vertex sets representing clades *C*(*G*) and clade splits *S*(*G*); see Fig. 1 for an example. For a clade *C* that is split into clades *C*_1_, *C*_2_, there is an edge from *C* to the clade split {*C*_1_, *C*_2_} in *G* as well as edges from {*C*_1_, *C*_2_} to *C*_1_ and *C*_2_. The root of *G* is the clade that represents the full taxon set *X* and the leaves of *G* are the taxa *𝓁* ∈ *X*. A CCD graph *represents (contains)* all trees that can be constructed by starting at the root clade and then recursively picking a clade split for each clade to progress. Second, for a clade split *S ∈* 𝒮 (*G*), let *π*(*S*) be the parent clade of *S*. Then the *conditional clade probability (CCP)* Pr_π_ (*S*) of *S* is the probability of picking *S* at *C*. So for each clade *C*, we have a distribution Pr_π_ on all clade splits 𝒮(*C*) of *C*. Now for a tree *T*, let 𝒮(*T*) be the clade splits of *T*. The probability of *T* in the CCD *D* is then given by the product of its clade splits’ CCPs [22]:

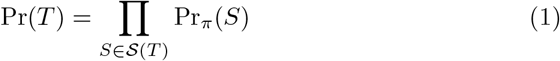

**Fig 1.**
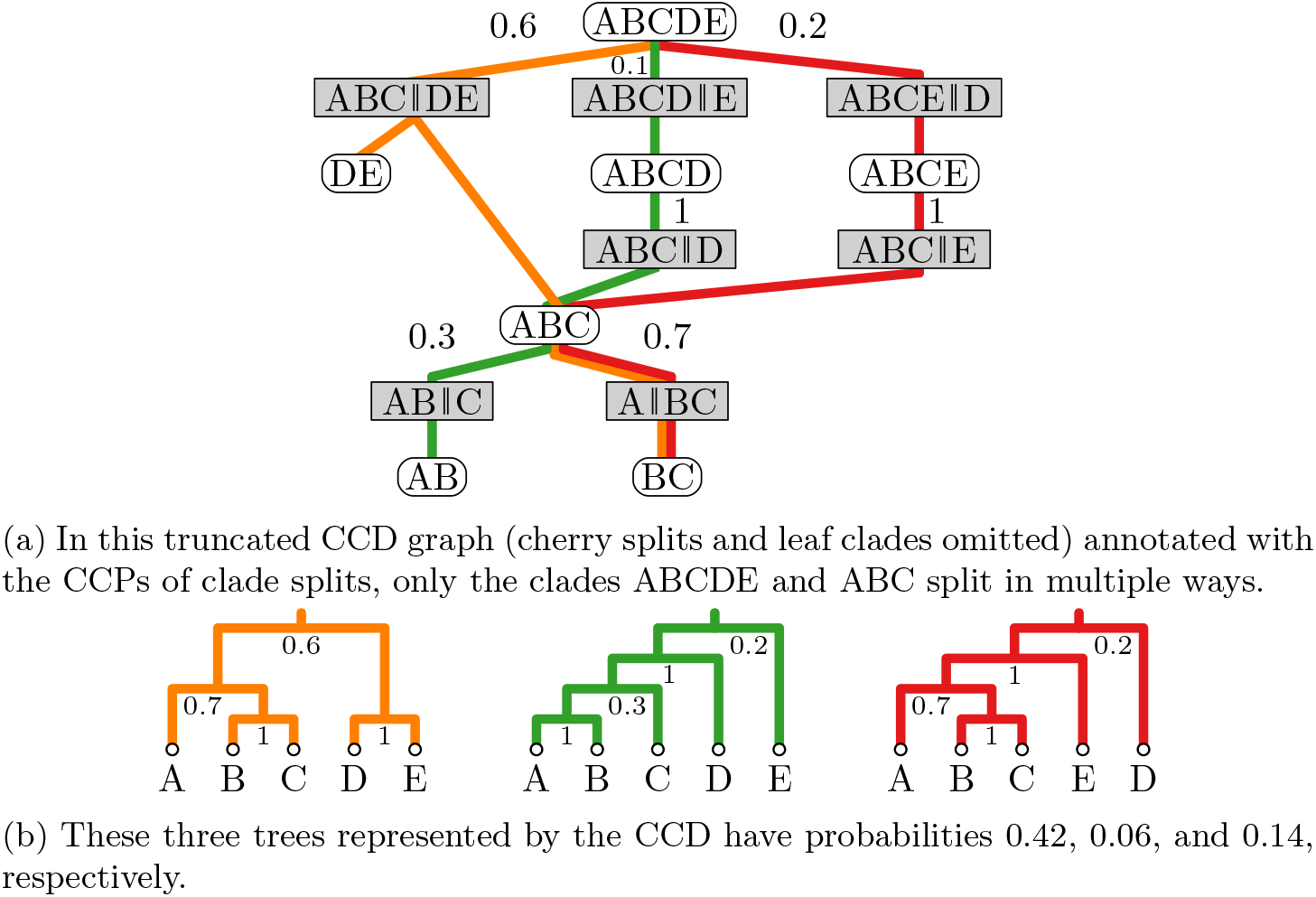
A CCD that contains six different trees.

For example, the first tree in Fig. 1b has probability 0.6 0.7 1 1. Larget [22] has shown that a CCD is a distribution on trees (that is, the probabilities of all trees in *G* add up to one). For a clade *C* and a clade split *S*, we define the probabilities Pr(*C*) and Pr(*S*) of *C* and *S* appearing in a random tree *T* in *D*, respectively. These probabilities can be computed for all clades and clade splits by a single pass over the CCD graph (cf. Berling et al. [3]). (We only need to compute the sum of probabilities of all paths from the root to this element where the probability of a path is given by the product of CCPs of its clade splits.)

(a) In this truncated CCD graph (cherry splits and leaf clades omitted) annotated with the CCPs of clade splits, only the clades ABCDE and ABC split in multiple ways.

(b) These three trees represented by the CCD have probabilities 0.42, 0.06, and 0.14, respectively.

The MAP tree of a CCD, that is, the tree with maximum probability, can be computed in linear time in the size of the CCD graph. We have recently shown that CCD MAP trees, in particular the CCD0 MAP tree, offer a point estimate that is as good or better than other methods such as MCC and greedy consensus trees in terms of accuracy and precision [3].

For further details and examples on CCDs, we refer to our recent paper [3] as well as Larget [22] and Lewis et al. [25].

#### CCD[*i*]

The parameters of a CCD0 are the non-zero clade probabilities and the CCD graph *G* contains all clades with non-zero probability as well as all clade splits that can be formed by them. More precisely, if *C*(*G*) contains clades *C, C*_1_, *C*_2_ with *C*_1_ ∪ *C*_2_ = *C* and *C*_1_ ∩ *C*_2_ = ∅, then the clade split {*C*_1_, *C*_2_} is in 𝒮(*G*). The clade probabilities can be converted into CCPs such that the probability of a tree is proportional to the product of its clades’ probabilities yet normalized [3].

The parameters of a CCD1 are the non-zero clade split probabilities. The CCD graph thus contains all clade splits with non-zero probability and the clades that form them. A CCD2 extends a CCD1 by having the probability of a clade split *S* being conditional not only on the parent π(*S*) but also on the sibling of π(*S*). This model is equivalent to the *subsplit directed acyclic graph (sDAG)* model by Matsen and colleagues [19, 38].

To populate the parameters of a CCD, we use the frequencies of clades and clade splits in a set of trees 𝒯, arising from an MCMC sample. Let *f* be the frequency of a clade/clade split appearing in 𝒯. Then for a clade *C* in a CCD0, the parameter of *C* is set to *f* (*C*)*/*|*T*| (i.e., equivalent to the Monte Carlo probability of *C*). In a CCD1, the CCP Pr_*π*_(*{C*_1_, *C*_2_*}*) of a clade split *{C*_1_, *C*_2_*}* of a clade *C* is set to *f* (*{C*_1_, *C*_2_*}*)*/f* (*C*), that is, the conditional probability of *C*_1_, *C*_2_ given *C*. In a CCD2, this is further distinguished by the sibling of *C*. The more complex CCD1 and CCD2 thus require more and more trees to estimate their parameters well. Other methods to populate these parameters are being studied [19, 39]. Furthermore, note that a CCD thus contain all trees that can amalgamated by the clades (CCD0) or clade splits (CCD1) observed in a sample.

#### Filtering a CCD

Let *D* be a CCD on taxa *X*, let *Y* ⊂ *X* be a subset of taxa, and let *Y* ^*c*^ = *X Y*. We are interested in the *skeleton (or reduced) CCD D* _*Y*_ of *D* on *Y*, that is, we want to filter out (marginalize out) the taxa *Y* ^*c*^. We may also call *D*|_*Y*_ the restriction of *D* to *Y*. If *Y* ^*c*^ corresponds to a clade *C*, we might write *D\C*. If we constructed *D* from a set of trees 𝒯, then we could obtain *D*|_*Y*_ by pruning each taxon in *Y* ^*c*^ from each tree in 𝒯 and reconstructing a CCD with the reduced trees. We now show how to obtain *D*|_*Y*_ from *D* efficiently. (It is not necessary to understand this for our rogue detection algorithm.)

Our algorithm to obtain *D*|_*Y*_ from *D* works recursively on the CCD graph *G* of *D* and handles a clade *C* only after all its child clades have been processed. A clade *C* with *C* ⊆ *Y* ^*c*^ simply gets removed. Otherwise, we construct or update the clade *E* of *D*|_*Y*_ to which *C* gets mapped and the corresponding clade splits including their parameters. If *D* was constructed from a tree set, then we only need to update clade frequencies for a CCD0, or clade and clade split frequencies for a CCD1, though we also describe the general case here.

We first describe how the CCD0 parameter (or the frequency) of a clade *E* in *D*|_*Y*_ is set. Let *ψ* : *C*(*D*) → *C*(*D*|_*Y*_) with *ψ*(*C*) → *C*∩*Y*, i.e., *ψ* maps each clade of *D* to its restriction in *D*|_*Y*_. Note that if *ψ*(*C*) = ∅, then *C* simply gets removed. Consider *ψ*^−1^(*E*) = {*C*_1_, …, *C*_*m*_}, the set of clades mapped to *E*. Note that *ψ*^−1^(*E*) can contain both a clade *C*_*i*_ and child clades of *C*_*i*_. Furthermore, no tree in *D* can contain two clades of *ψ*^−1^(*E*) that do not have an ancestor-descendant relationship. Hence, each *C*_*i*_, *i* ∈ [*m*], contributes to the parameter of *E* with its own parameter (i.e., the frequencies add up) except for the proportion where also a child of *C*_*i*_ is in *ψ*^−1^(*E*) (i.e., where two clades of one tree are mapped to *E*). This proportion derives directly from the CCPs of *C*’s clade splits as we know with what probability we have *C*_*i*_ and a child of *C*_*i*_ together. Thus the algorithm updates the current value of *E* when processing *C*. For a CCD0, once we have processed the root clade, we again convert the clade parameters into the CCPs of all clade splits in *D*|_*Y*_ (as in a new CCD0 [3]).

Next, we describe how to update the parameter (frequency or CCP) of a clade split *R* in *D*|_*Y*_ for a CCD1. (Recall that for a CCD0, no information on clade splits is needed.) Let *ϕ*^−1^(*R*) = {*S*_1_, …, *S*_*𝓁*_}, the set of clade splits mapped to *R*. Note that no two clade splits in *ϕ*^−1^(*R*) can have an ancestor-descendant relationship, i.e., lie on the same path from the root to a leaf. Hence, the frequency of *R* equals the sum of frequencies of clade splits in *ϕ*^−1^(*R*). We can also directly compute Pr_*π*_(*R*), the CCP of *R*. Let *E* = *π*(*R*) (the parent of *R*) and *C*_*i*_ = *π*(*S*_*i*_), *i ∈* [*𝓁*]. For each *S*_*i*_, the contribution of Pr_*π*_(*S*_*i*_) to Pr_*π*_(*R*) is equivalent to the contribution of *C*_*i*_ to *E*, which is given by Pr(*C*_*i*_)*/* Pr(*E*). Therefore, 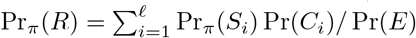. Note that Pr(*E*) only acts as normalizing constant, which we can compute in a second pass over all clades.

Overall, we can compute *D*|_*Y*_ in linear time in the size of *G*(*D*), so in the number of clades and clade splits in *D*.

### 2.2 Phylogenetic Posterior Entropy

Entropy is a measure of uncertainty or randomness of a distribution. More precisely, the *information content* of an event *x* is defined as − log Pr(*x*)^2^. The *entropy* is then the expected amount of information (or “surprise”) in a distribution and a high entropy thus indicates high uncertainty. Furthermore, the entropy is comparable among datasets and does not depend on a particular metric space. For a (posterior) tree distribution *D*, the *phylogenetic (posterior) entropy* of *D* is given by the following formula:

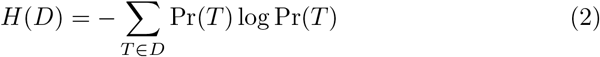

Given the intractably large number of trees in a CCD *D* for nontrivial problems, we cannot compute *H*(*D*) directly with Eq. (2). However, we can simplify this as follows. First, consider only one tree *T* and use Eq. (1):

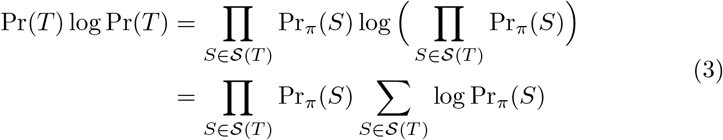

Next consider the term log Pr_*π*_(*S*) for a particular clade split *S*. Note that when summing over all trees in Eq. (2), the term appears once for each tree *T* ^*′*^ that contains *S* and with Pr(*T* ^*′*^) as weight. Summing over all trees, we thus get a total weight of Pr(*S*), i.e., the probability of *S* appearing in a tree in *D*. Summing over all clade splits, we get the following simple formula for the phylogenetic entropy of *D*:

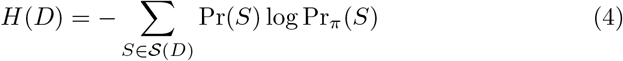

Hence, we can compute the entropy of *D* in one pass over the CCD graph with the explicit formula Eq. (4) or, alternatively, with the recursive formula by Lewis et al. [25].

The *number equivalent* ^3^of an entropy *H* is the number of equally likely elements needed to get *H* and is thus given by exp(*H*). For example, if we consider the set of all four-taxa trees and give them equal probability (e.g. 1*/*15), then their distribution *D* has entropy *H*(*D*) ≈ 2.708 and exp(*H*(*D*)) = 15.

### 2.3 Rogue Scores

We now define rogue scoring functions for a set of taxa *C* (also called *dropset*, which can be a single taxon, non-trivial clade, or set of clades) with respect to a posterior tree distribution *D* (here represented by a CCD). The main idea is to measure the difference in entropy between *C* being included and excluded from *D*.

The first scoring function measures the total reduction in entropy by excluding all taxa in *C*. More precisely, with *Y* = *X\C*, we define the *total rogue score w*(*C*) of *C*:

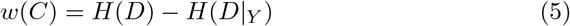

We might also consider the *proportional total rogue score* 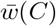 by dividing by the size of *C*:

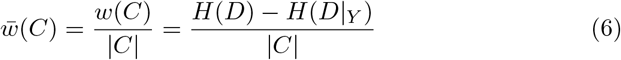

For a leaf *𝓁*, we can interpret the number equivalent exp(*w*(*𝓁*)) of *w*(*𝓁*) as an estimate of the number of equally likely placements of *𝓁* to a tree to obtain that entropy. For example, if we construct a distribution *D* by attaching to a three-taxa tree a fourth leaf *𝓁*_4_ thrice and each time at a different edge, we get *H*(*D*) = *w*(*𝓁*_4_) ≈ 1.099 and exp(*w*(*𝓁*_4_)) = 3. For a non-leaf clade *C* on the other hand, the total rogue score includes the variance caused by *C* itself as well as all subclades and taxa of *C*. For example, a clade *C* can be placed with 100% support, but still have high uncertainty within, and thus a high score *w*(*C*).

To capture the variance caused by the placement of a clade *C*, we define the *clade rogue score ŵ* (*C*) as the reduction in entropy by excluding *C* but not the variance within *C*:

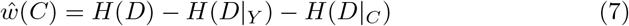

We can interpret the number equivalent exp (*ŵ*(*C*)) of *w*(*C*) as an estimate of the number of equally likely placements of *C*. The clade rogue score is particularly of interest for clade with high support, for example those in a summary tree. For a leaf *𝓁*, note that 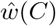.

Concerning the computation of *w*(*C*), *ŵ*(*C*) and with Eq. (4) in mind, note that removing a clade can change the entropy in a nontrivial way: While all removed clade splits containing *C* directly contribute to *w*(*C*), both the probability and CCP of other clade splits can change as well. Hence, we construct^4^*D*|_*Y*_ to compute *H*(*D*|_*Y*_). This takes 𝒪(|*D*|) time per clade and thus 𝒪 (|*D*|^2^) for all clades. In practice, we expect to have a maximum size *k* for clades considered.

An alternative score *w*^*′*^(*C*) for a clade *C* is to sum the entropy contribution of all clade splits that contain *C*:

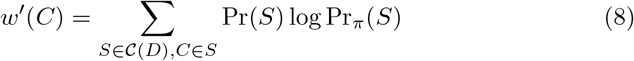

This score can be computed fast (namely, in linear time for all clades) and comes close to the one used by Smith [31], but it does not capture the whole reduction in variance in the CCD when dropping *C*.

### 2.4 Analyses

Next we describe how rogue scores can be used in a posterior analysis to (i) identify candidates for true rogues, (ii) assess relative uncertainty of clade placements, and (iii) find the skeleton in the forest.

#### Rogue Analysis

We start with finding rogue candidates and visualizing relative uncertainty. For a given CCD *D*, a maximum clade size *k*, and a threshold *t*, we compute the clade rogue score for each clade in *D* of size at most *k* and support at least *t*, and can then visualise and analyse the outcome as follows.

First, using a ranked plot of the scores allows for a visual comparison of individual clades. This approach can help identify outliers (i.e., potential true rogues), reveal trends, or indicate whether entropy is evenly distributed. Here we suggest a probability threshold *t* of 10% and *k* = 5. Examples of this are provided in Section 3.

Second, we can annotate a summary tree (in our case, a CCD MAP tree [3]) with the clade rogue score as meta data for each clade. Using standard tree visualisation tools, the scores can be depicted by node size or edge color. This can offer a nuanced understanding of which clades contribute to the variance and low support values in the tree, supplementing the information that clade support provides. The tool RogueAnalysis, available through the CCD package, outputs a file with rogue scores for each clade, as well as an annotated CCD MAP tree.

#### Skeleton Analysis

A more comprehensive analysis of rogues in a distribution aims at finding a *skeleton distribution* that maintains essential phylogenetic signals while minimizing the impact of noisy, hard-to-place clades. This skeleton distribution can then be used for more accurate downstream analyses, providing a clearer understanding of the evolutionary relationships among the taxa. The basic idea is to iteratively remove rogue clades until we arrive at a skeleton CCD that satisfies some criterion. Example criteria are a threshold on the remaining entropy *H*(*D*|_*Y*_) and a threshold for the probability of each clade in the MAP tree of *D*|_*Y*_. We further assume that we have a maximum size *k* and a probability threshold *t* for a clade we might remove.

Our algorithm to compute a skeleton CCD for some criterion and maximum clade size *k* is a greedy dynamic program: Let *D*_0_ be our initial CCD distribution on taxon set *X* and *n* = |*X*|. For *i* ∈ [*n*], the algorithm store as *D*_*i*_ a CCD on *n* − *i* taxa that has the lowest entropy among all considered. More precisely, for all *j* ≤ *k* and each clade *C* of size *j* in *D*_*i*−*j*_ (and probability at least *t*), the algorithm computes *H*(*D*_*i*−*j*_ *C*) and sets *D*_*i*_ = *D*_*i*−*j*_ *C* if this is a new lowest entropy. (Equivalently, we pick *C* with the highest total rogue score that results in *n*− *i* taxa.) It stops once the stopping criterion is fulfilled, the entropy *H*(*D*_*i*_) is zero, or if for the last *k* rounds no improvement has been made. In the latter case, all clades of size at most *k* are uniquely placed. The algorithm has a dynamic program component as it considers multiple previous solutions but overall greedily decides on a sequence of clades to remove. There can theoretically be a sequence of clades whose removal would result in a lower entropy with *n − i* remaining taxa, but that would not have been optimal at intermediate steps. We demonstrate a skeleton analysis in Section 3.

Each round of the algorithm takes 𝒪 (|*D*| ^2^) time (and 𝒪(*n* |*D*|) for *k* = 1). For very noisy datasets, there can a significant number of small clades with very low probability. Instead of considering such clades potential rogue clades, it makes more sense to remove their taxa one by one. Hence, to speed up the algorithm tremendously, we added the probability threshold *t* for a clade *C* to the algorithm. For example, such a threshold could be 10%.

The parameters *k* and *t* as well as the stopping criteria are parameters for the tool SkeletonAnalysis, available through the CCD package. This tool outputs a list of rogue taxa removed to reach the skeleton CCD as well as the CCD MAP tree annotated with rogue placements (described next). Furthermore, it can create a copy of the given tree set reduced to the taxon set of the skeleton CCD for further analysis.

#### Placement Analysis

In addition to a rogue and skeleton analysis, we can also compute and visualize all possible placements of a clade *C* in a tree *T* of a reduced CCD *D\C*. In particular, this is of interest for clades *C* with relative high rogue score and *T* being the CCD MAP tree of *D\C*. To this end, we compute for each parent clade split *S* of *C* in *D* whether *S* is mapped to an edge *e* of *T* in *D\C*. If so, we store for *e* the probability of *S* being on a path from the root to *C*. Note that multiple splits can be mapped to *e*. On the other hand, not all splits are mapped to edges of *T* and there can thus be “lost” probability of placements of *C* with respect to *T*. Hence we recommend a placement analysis only for cases where *T* has a high probability, e.g., the MAP tree of a skeleton CCD. The tool SkeletonAnalysis annotates the MAP tree with rogue placement metadata for all clades removed in obtaining the skeleton CCD.

### 2.5 Datasets

We selected the following sets of posterior tree samples inferred using BEAST2 [5] to demonstrate our analyses across a range of datasets varying in size (but not too large to display trees) and entropy:

DS7 A set of 10,000 trees on 59 Malagasy lemur taxa from multiple gene loci [37], used as dataset DS7 in a suite of popular benchmarking datasets in phylogenetics [21] with an entropy around 10.

SCC A set of 689 single-cell cancer (SCC) trees on 86 taxa from Chen et al. [8], who used SNV data [20] from single colorectal cancer patient [24]. This distribution has an entropy of around 69.

RSV2 A set of 50,000 trees of 129 time-stamped molecular sequence coding for RSVA’s glycoprotein [40, 41] following the model (and tutorial) of LinguaPhylo [10], which has an entropy of around 45.

In addition, we used LinguaPhylo [10] to simulate 100 datasets under a simple Yule model with 100 taxa and a sequence length of 300 sites; for more details see Berling et al. [3] with the only difference being that we used a non-fixed birth-death prior (log-normal distribution). For each simulation, we obtained two replicates of 10,000 trees after burnin using BEAST2 [5]. We call this dataset Yule100.

## 3 Results

In this section, we demonstrate how our algorithms and tools can be applied in a phylogenetic analysis. Our goal is to showcase the practical use of these methods on four case studies, while outlining some of its limitations. The examples provided illustrate how the rogue and skeleton analysis can help identify rogue taxa, assess clade uncertainty, and highlight sources of variance in tree sets. We anticipate that practitioners will adapt and refine these methods for their specific datasets, potentially uncovering unique insights and applications beyond what is shown here. We also conducted a small experiment on whether skeletonization introduces a systematic bias. Throughout, we used the CCD0 model for estimates of the posterior distribution, MAP trees, and phylogenetic entropy calculations.

### Demonstrating Rogue Analysis

To illustrate the practical utility of a rogue analysis, we analyzed the three real datasets (Section 2.5). First, we generated rank plots of the clade rogue scores for clades up to a size of 5 and with at least 10% support. Second, we annotated a CCD0 MAP tree from each dataset with clade rogue scores and visualized the three trees where the edge color represents the scores using FigTree [29]. See Fig. 2 for all plots and trees. In interpreting these figures, it should be note that these should be considered snapshots during an investigative analysis and not as final results meant for print.

**Fig 2.**
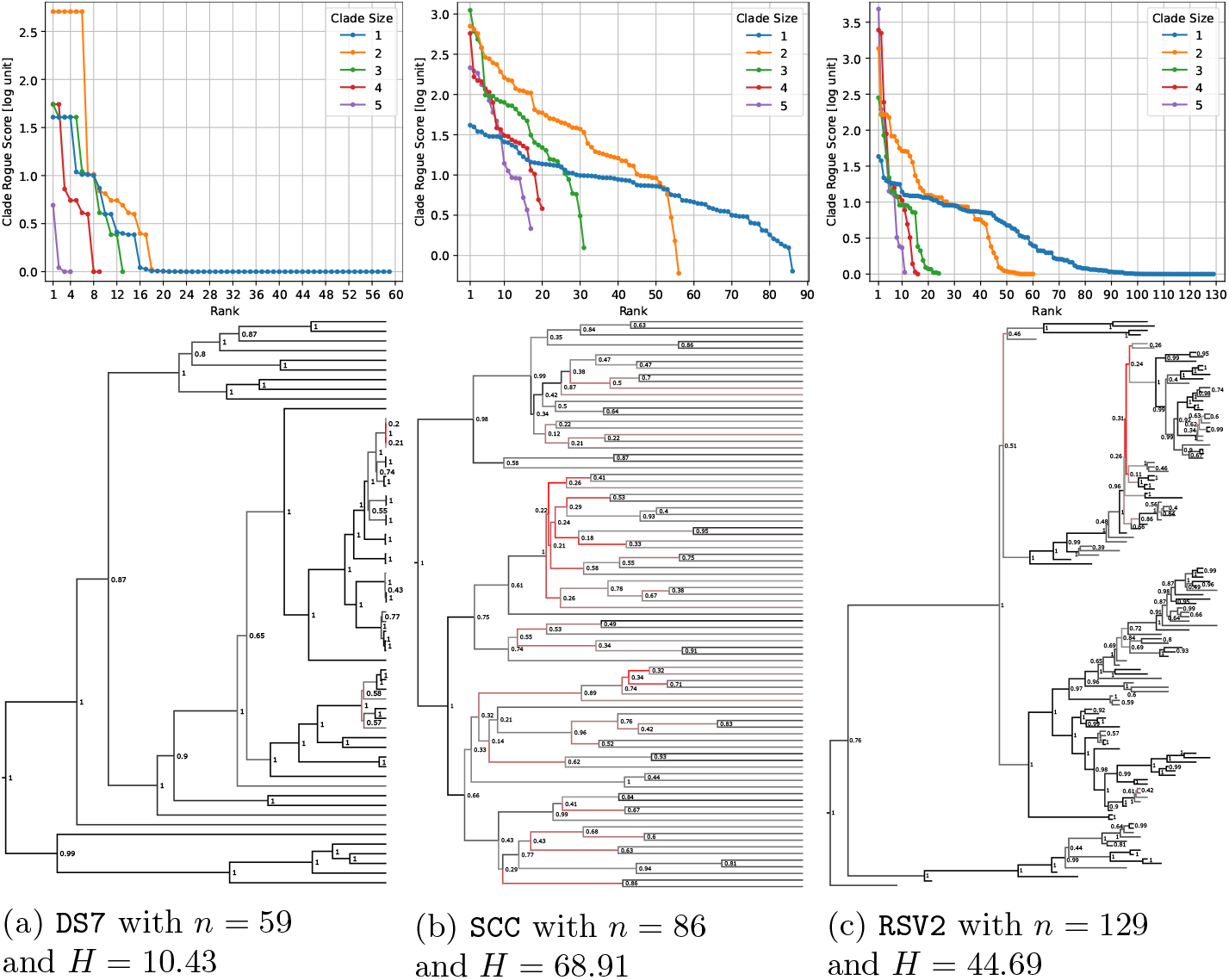
Rogue analysis for three datasets with the rank plots at the top for clades of size up to 5 and with at least 10% support (clades of a certain size are ranked based on their clade rogue score, higher score indicating less well placed). The CCD0 MAP trees at the bottom are annotated with clade supports in text and with clade rogues scores in color from black (low) via grey to red (high).

Taking a closer look at DS7 in Fig. 2a first, we find in the rank plot that there are four taxa with a significantly higher entropy score than all others. Investigating the MSA reveals that the corresponding sequences are near-identical thus making relative placements near impossible. This can also be observed in the MAP tree, where their clade has nearly zero height. Furthermore, the six cherries (clades of size two) and four clades of size three with high rogue scores correspond to the different clades formed by these taxa. Note that not all clades present in the rank plot also appear as clades in the MAP tree.

Next, consider SCC in Fig. 2b. We observe that there are no taxa with significant higher entropy than others, but a relative trend from less-well to well placed taxa; one taxon stands out though with a negative rogue score indication a stabilizing contribution of this taxa, which is possible due to the non-linear nature of the entropy function. In general, we observe that many cherries are less well placed than taxa, which also becomes apparent in the annotated MAP tree. Furthermore, we can see in the MAP tree that at some clades with low support, the two child clades are differently well placed (as the different rogue scores are highlighted by different colours). We thus get a more nuanced picture of the uncertainty. The situation is similar for RSV2 in Fig. 2c. We see that several large clades have significant higher rogue scores than all taxa, with many taxa being perfectly placed.

### Demonstrating Skeleton Analysis

Next we conducted a skeleton analysis on the RSV2 and SCC datasets with our algorithm to repeatedly remove the clade with the highest total rogue score to obtain skeleton CCDs; see Figs. 3 and 4. We set 10 as the maximum rogue clade size, only considered clades with at least 50% probability, and, as stopping criteria, used thresholds on the entropy of 10 and 5. We are showing both cloudograms from DensiTree [6] and CCD0 MAP trees for the resulting three datasets each (full, *H* ≤ 10, and *H* ≤ 5). The number of taxa got reduced, for RSV2, from 129 via 94 to 84 with a maximum rogue size of 3 (once and three cherries); for SCC the number of taxa went from 86 via 33 to 25 and the largest rogue had size 2 (eleven times). Unearthing the skeleton took 5 seconds for RSV2 and 13s seconds for SCC, which for the former was faster than parsing the trees.

**Fig 3.**
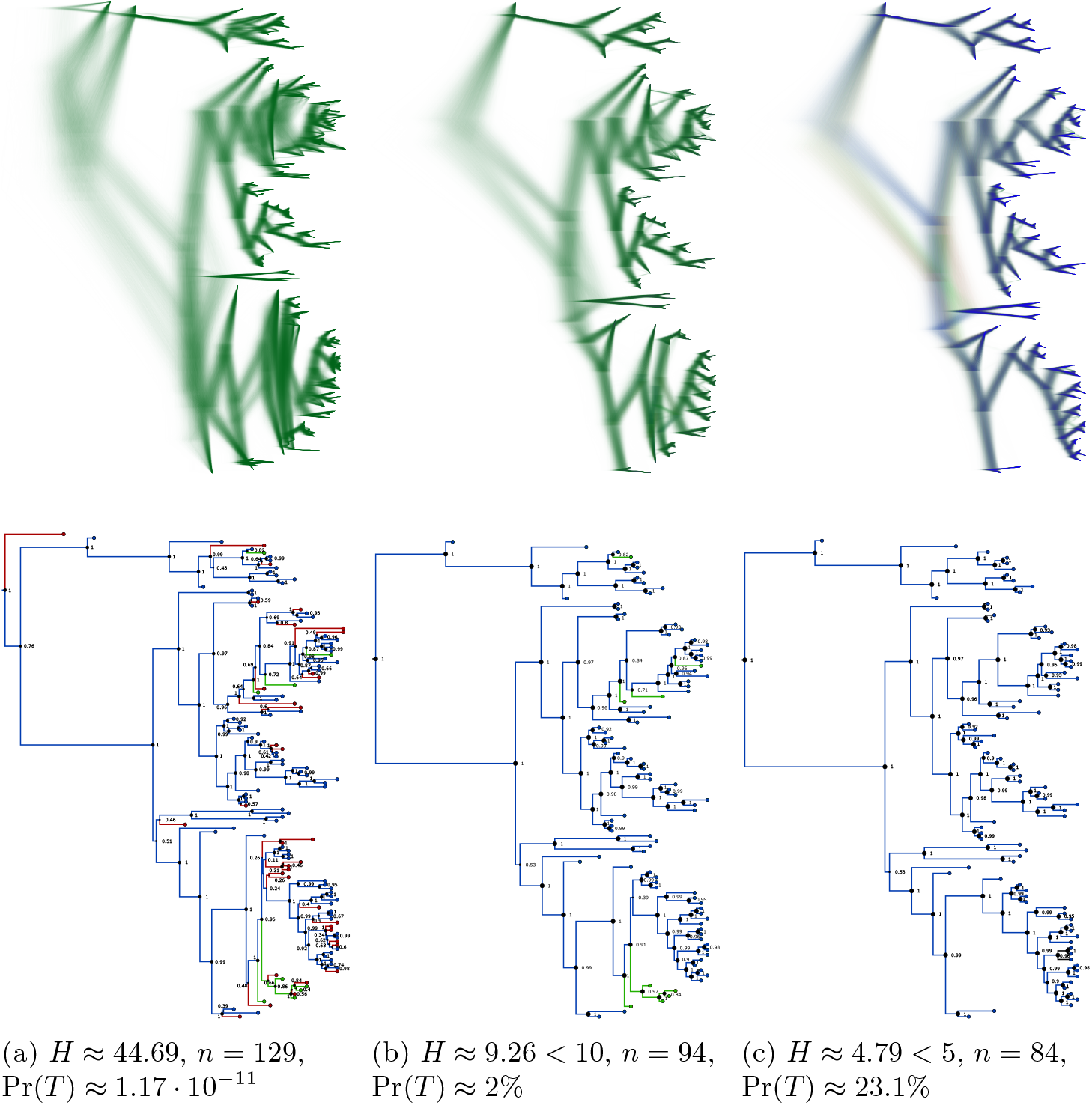
Skeleton analysis for the RSV2 dataset with entropy thresholds 10 and 5 showing cloudograms of the reduced tree sets and the MAP tree *T* of the skeleton CCD where *n* is the number of (remaining) taxa. In the cloudograms, the most common tree (topology) is blue, the second one red, while all others are green. In the point estimates, the numbers show the clade supports and the subtree on the taxa of the skeleton with *H* ≤ 5 is coloured blue, additional edges on the taxa also in the skeleton with *H* ≤ 10 green, and those only on the full taxon set red.

**Fig 4.**
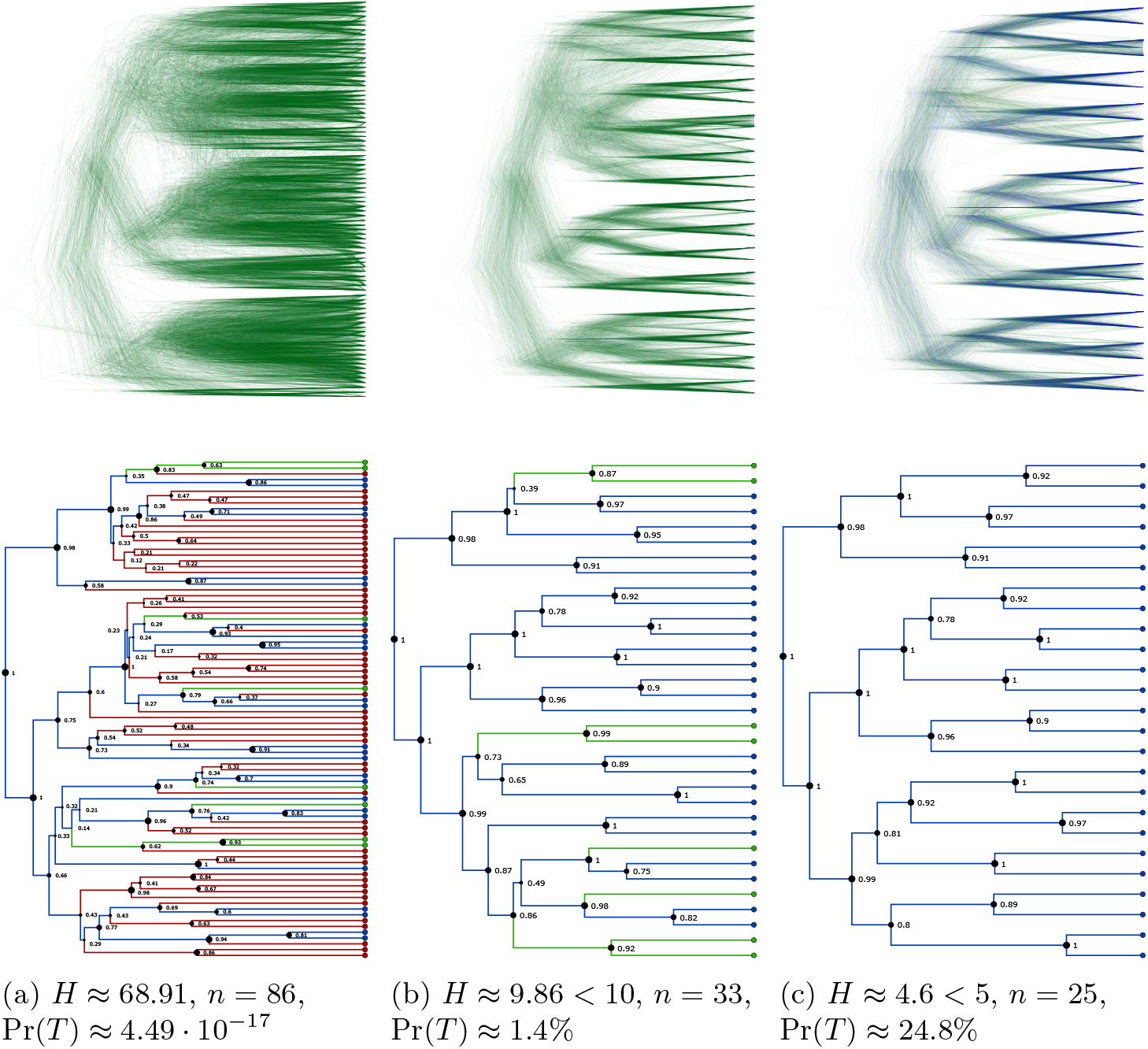
Skeleton analysis for the single-cell cancer dataset SCC with entropy thresholds 10 and 5 showing cloudograms of the reduced tree sets and the MAP tree *T* of the skeleton CCD0 where *n* is the number of (remaining) taxa. In the cloudograms, the most common tree (topology) is blue, the second one red, while all others are green. In the point estimates, the numbers show the clade supports and the subtree on the taxa of the skeleton with *H* ≤ 5 is coloured blue, additional edges on the taxa also in the skeleton with *H* ≤ 10 green, and those only on the full taxon set red.

The cloudograms show the progression from a very noisy full dataset to stronger and stronger signal, where most of the noise comes from the different height estimates and not different trees. DensiTree colours the most common tree (topology) blue, which becomes dominant for *H*≤ 5.

In the CCD0 MAP trees, we see how the support of all clades drastically increases, e.g. for SCC the lowest value increasing from 14% to 39% for *H* ≤ 10 and finally to 78% for *H* ≤ 5. (Recall that the support values are the main criteria for most ML rogue detection algorithms.) Furthermore, the probabilities of the CCD0 MAP tree increase from 4.49 · 10^−17^ to 0.248 for SCC and from 1.17 · 10^−11^ to 0.231 for RSV2. This shows that, say for RSV2, the noisy distribution on 129 taxa contained a skeleton distribution on still 84 taxa (65%) with very strong signal, namely over 23% being one tree.

### Demonstrating Placement Analysis

We can further analyse how each removed clade *C* relates to the previously obtained skeleton CCD with *H* ≤ 5. In particular, using the CCD0 MAP tree *T* (with 23.1% probability), we compute on which edges of *T* the clade *C* would be placed on. Fig. 5 shows the different placements for two taxa for the CCD0 MAP tree of the RSV2 skeleton. In practice, this would be part of an interactive analysis. Note that while the placement of *C* in the skeleton might be relatively certain, many different rogues attach to the same edges, which caused the initial high entropy. On the other hand, we can compute the expected number of removed clades to be placed on any edge of *T*.

**Fig 5.**
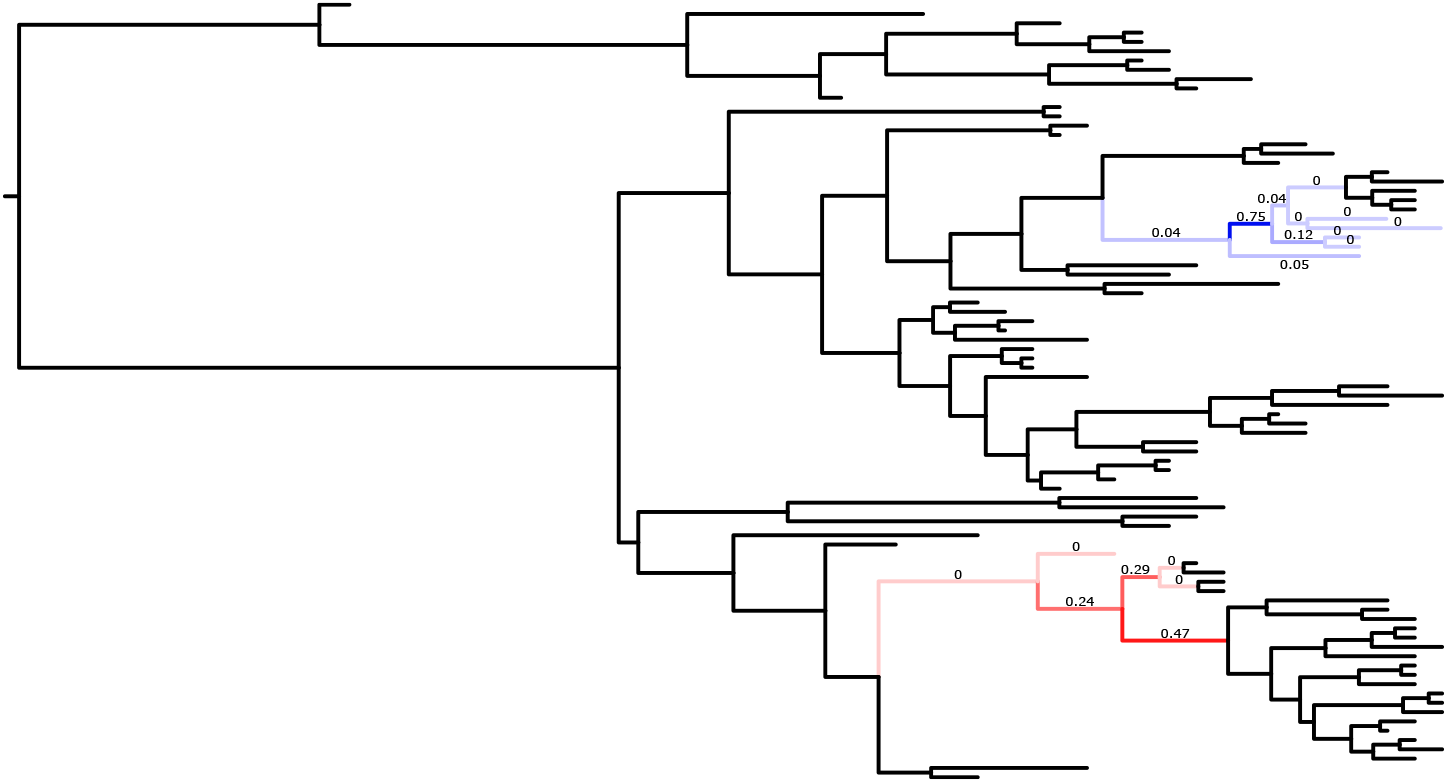
On the CCD0 MAP tree of the RSV2 skeleton with *H* ≤ 5, visualisation of the different placements of the 1st removed taxon (red) and the 28th removed taxon (blue) with color and label indicating the proportion (with 2 digit precision, resulting in displayed 0s).

We want to remark that for this analysis to be effective, we require a reference tree *T* with high absolute probability since otherwise many placements of a clade may not be on edges of *T*. For example, if instead we use the CCD0 MAP tree of the RSV2 skeleton distribution for *H* ≤ 10, which has a probability of about 2%, we find that only 46% of placements of the 1st removed taxon is on *T* ; for *H* ≤ 5, it is nearly 100%.

### Well-Calibrated Simulation Study

To test whether constructing a skeleton with our entropy-based algorithm introduces a bias on any of the evolutionary model parameters, we conducted a well-calibrated simulation study on Yule100 [26]. More precisely, with the same parameters as above, we ran the algorithm to obtain a skeleton CCD for *H* ≤ 5 and 10 for both replicates of each of the one hundred simulations. For *j* ∈ {5, 10}, let *Y*_*j*_ be the remaining taxa of the skeleton with *H* ≤ *j*. Firstly we reduced the original tree set (posterior sample) to *Y*_*j*_; let us call this the *reduced tree set* on *Y*_*j*_. Secondly we rerun the phylogenetic inference with BEAST2 on the sequences corresponding to *Y*_*j*_; let us call the resulting posterior sample the *rerun tree set* on *Y*_*j*_. For the reduced tree sets, we compared the tree height and the tree length to true value of the simulation; for the rerun tree sets, we also compared the birth rate, kappa, shape parameter, and base frequencies. (Note that for the reduced tree sets, the estimates of these parameters equal those on the full tree set.) Overall, we could not detect a bias for any of the parameters and the studies passed the coverage test. Coverage details are given in Table 1.

**Table 1.** Coverage (in %) for Yule100 in full and its skeleton distributions (with entropy *H* below 10 and 5) obtained by removing taxa from the trees (reduced) as well as by rerunning the analysis on the remaining sequences (rerun).

| Parameter | Full | $H \leq 10$ | | $H \leq 5$ | |
| --- | --- | --- | --- | --- | --- |
|  |  | Reduced | Rerun | Reduced | Rerun |
| tree height | 94 | 94 | 94 | 94 | 94 |
| tree length | 92 | 95 | 92 | 95 | 95 |
| birth rate | 93 | - | 94 | - | 94 |
| kappa | 94 | - | 93 | - | 91 |
| shape | 97 | - | 96 | - | 93 |
| frequencies (avg) | 95.5 | - | 95.5 | - | 97.5 |

The initial entropy of the CCD0s was 30.54 on average. The 100 taxa was reduced by an average of 22.79 and 34.67 taxa for the thresholds of 10 and 5, respectively. Furthermore, comparing the two replicates per simulation, the symmetric difference of the sets of removed rogues was on average 3.12 and 4.3 taxa for the thresholds of 10 and 5, respectively. This shows that CCDs and the skeleton algorithm are stable.

### Rerunning RSV2

In addition to the well-calibrated simulation study, we also rerun the inference for RSV2 on reduced MSAs based on the taxon sets of the reduced skeleton CCDs. The resulting CCD0s had very similar entropies of 9.32 (cf. 9.26 of the reduced skeleton CCD) for *H* ≤ 10 and 4.9 (cf. 4.79) for *H* ≤ 5; the CCD0 MAP tree were the same with slightly increased probabilities of 3.3% (cf. reduced 2%) on *Y*_10_ and 31.6% (cf. reduced 23.1%) on *Y*_5_.

### Comparison with RogueRaNok

Performing rogue detection on the example data set that comes with RogueRaNok^5^ results in 28 removals using RogueRaNok. With CCD1 and a removal limit of 28, we find that 19 taxa identified for removal are identical while 9 differ. The entropy of the tree set with suggested rogues removed is 49.11 for RogueNaRok’s selection and 45.99 for the CCD based one. This demonstrates the optimisation criterion used for detecting rogues can have a substantial impact on the resulting tree set.

## 4 Discussion

In this discussion, we address two key topics: the newly introduced applications for posterior analysis and the potential for systematic errors introduced by the removal of taxa.

Our main application of the phylogenetic entropy-based rogue detection is the unearthing of the skeleton tree distribution, that is, the remaining distribution after repeatedly removing the clade with the highest rogue score. The algorithm is designed to be both efficient and minimally invasive, removing the smallest number of taxa to reveal a strong phylogenetic signal. As demonstrated in our analysis of real datasets, this approach can uncover clear phylogenetic relationship within noisy tree distributions. Furthermore, the resulting summary tree has a high probability and exhibits high clade support values. Alternatively, the gained information can be used to rerun the inference analysis on a reduced set of sequences.

Since the algorithm is based on information theory and requires no choice of a tree metric (or geometric treespace), and there is no additional cost for analysing larger clades, it can deal with various types of datasets. While for the RSV2 dataset mostly single taxa were removed and very strong signal was revealed with still 63% of the taxa, less than a third was kept for SCC dataset and an entropy below five. There the number of removed taxa was amplified by good pairings of taxa into cherries which in turn were badly placed, resulting in the removal of eleven cherries.

In addition to skeletonization, we introduced new methods to analyze the placement of specific taxa and clades within the tree distribution. This includes evaluating clade placements in relation to the skeleton summary tree and utilizing the entropy-based clade rogue score to assess uncertainty within the posterior distribution. As demonstrated in our examples, this approach can effectively identify outliers or potential rogue taxa, uncover underlying trends, and offer a more detailed understanding of clade placements that goes beyond the insights provided by traditional clade support values and cloudograms.

These applications may be particularly valuable to systematists who, for various reasons, cannot or prefer not to exclude any taxa from their analyses. By investigating the uncertain placements of specific taxa using the clade rogue score, they can still gain insights into the stability and reliability of their phylogenetic inferences.

It is important to note that the marginal distribution of trees obtained by removing rogue taxa is not necessarily equivalent to the distribution obtained by rerunning an analysis on an alignment from which rogue taxa have been excluded [4]. One reason is that rogue taxa impact parameter estimates of the site and clock models, which may impact the timing and topology of the trees. Another reason is that rogue taxa can repel taxa from certain clades when included, but not when excluded, resulting in different clade distributions. Both of these effects have been described elsewhere [4]. In general, the tree set with rogue taxa pruned is based on more data (than the result of re-running on the taxa subset), and therefore may be assumed to be more reliable. However that is predicated on the rogues not actually being statistical outliers. If the removed rogues are in fact outliers that are not well-described by the phylogenetic model, then re-running without them may actually produce better estimates. Rerunning the analysis on the alignment without rogue taxa can also be useful to identify how much information is obtained from the rogue taxa. In our analysis of the RSV2 dataset, for example, we found that the contribution of rogue taxa was negligible.

Furthermore, our well-calibrated simulation study showed that computing skeleton CCDs did not introduce a systematic bias under the simple Yule model. This was the case for both removing rogues directly from the trees as well as rerunning the analysis on the remaining taxa. It remains to be demonstrated how well these results generalise to other systems and datasets. Further research is needed to explore the effects of removing large numbers of taxa, whether directly from the CCDs or from the sequences, when selected based on phylogenetic entropy. In particular, this could be studied for models of datasets that are often very noisy, such as single-cell sequencing data [8], as well as for other real datasets.

For all experiments, we used the CCD0 model. This choice is supported by findings by Berling et al. [3], which showed that for non-trivial (simulated) datasets the regimes where CCD1 and CCD2 (or even just the tree sample) offer better models can only be reached, if at all, by a huge number of sampled trees. Further studies into alternative methods to populate the parameters of CCDs (cf. Zhang and Matsen [38, 39], Jun et al. [19]) as well as into model selection for real datasets may lead to the adoption of other models and even better results.

## 5 Conclusion

In this paper, we introduced new rogue detection methods for Bayesian posterior tree distributions based on phylogenetic entropy to address the challenges posed by noisy data. Our approaches enable new applications in posterior analyses, particularly for uncovering skeleton distributions and conducting fine-grained analysis of of uncertainty in clade placement. These methods are implemented in the CCD package and integrated into DensiTree.

We have demonstrated several analyses and visualizations that utilize entropy-based rogue detection, offering a foundation for further exploration. Researchers and systematists with specific datasets and hypotheses may discover their own ways to apply these algorithms and rogue scores to enhance their analyses. For example, it would be interesting to see whether rogue scores can help detect hybrids and other signals that conflict with a tree-like histories. In preliminary work on a particularly noisy dataset where the Bayesian MCMC algorithm did not converge, rogue detection facilitated the identification of a subset of taxa that allowed successful convergence (data not shown).

Future work could extend these methods, for example, by improving visualization tools, and by exploring rogue detection linked over multiple tree distributions. The impact of entropy-based rogue removal on specific datasets also remains an intriguing avenue for further research.

## Acknowledgements

JK and AJD were partially supported by the Beyond Prediction Data Science Research Programme (MBIE grant UOAX1932). We would like to thank Caroline Puente-Lelievre and Jordan Douglas for insightful discussions as well as Kylie Chen and Walter Xie for technical help concerning the datasets.

## Footnotes

1 GitHub CCD package github.com/CompEvol/CCD

2 Throughout, we use the natural logarithm.

3 It is sometimes also called “effective number of elements” and “number of states” [18].

4 In fact, to be more efficient, we can construct a partially new CCD that reuses the parts of *D* that remain unchanged.

5 Available from https://github.com/aberer/RogueNaRok/blob/master/example/150.bs.

